# YDS-GlueFold: Surpassing AlphaFold 3-Type Models for Molecular Glue-Induced Ternary Complex Prediction

**DOI:** 10.1101/2024.12.23.630090

**Authors:** Xing Che, The Statistical Mechanics Inference Team

## Abstract

Molecular glues represent an innovative class of drugs that enable previously impossible protein-protein interactions, but their rational design remains challenging, a problem that accurate ternary complex modeling can significantly address. Here we present YDS-GlueFold, a novel computational approach that enhances AlphaFold 3-type models by incorporating guided diffusion during inference to accurately predict molecular glue-induced ternary complex structures. We demonstrate YDS-GlueFold’s capabilities across eight diverse test cases, including both E3 ligase-based systems (VHL, CRBN complexes with mTOR-FRB, NEK7, and VAV1-SH3c, and KBTBD4 complexes with HDAC1) and non-E3 ligase complexes (FKBP12 complexes with mTOR-FRB, BRD9 and QDPR). The model achieves remarkable accuracy with RMSD values consistently below 2.5 Å compared to experimental structures. Importantly, 7 out of 8 test cases involve protein-protein pairs that were not present in the AlphaFold 3 training set, providing a rigorous test of the model’s ability to generalize beyond its training data. Notably, in the FKBP12 case, YDS-GlueFold correctly predicts a novel interface configuration instead of defaulting to known interactions present in training data, further demonstrating true generalization rather than mere memorization. Our results suggest that strategic enhancement of the inference process through guided diffusion can significantly improve ternary complex prediction accuracy, potentially accelerating the development of molecular glue therapeutics for previously undruggable targets.

## Introduction

Drug discovery has traditionally been limited by our ability to target only proteins with well-defined binding pockets suitable for small molecules. This constraint has left many disease-relevant proteins, particularly those operating through protein-protein interactions, beyond therapeutic reach. However, molecular glues—an innovative class of drugs—are transforming this paradigm by acting as molecular matchmakers that enable previously impossible protein-protein interactions^1^.

Molecular glues operate through different mechanisms of action. Classical molecular glues, exemplified by rapamycin, act as molecular “adhesives” by stabilizing specific protein-protein interactions that would not occur spontaneously, such as the FKBP12-mTOR complex that leads to mTOR inhibition. Another important class of molecular glues triggers protein degradation, harnessing cellular protein degradation machinery by facilitating ternary complex formation between an E3 ubiquitin ligase (such as VHL or CRBN) and a target protein. These molecular recognition events involve complex cooperative binding, where the molecular glue often induces conformational changes in one protein that create new interaction surfaces to attract another protein. While this intricate interplay enables molecular glues to modulate previously “undruggable” proteins, it also makes their rational design particularly challenging^2^.

Computational modeling is a common strategy for exploring rational drug design. However, traditional methods, originally developed for binary interactions such as ligand-protein or protein-protein complexes, face significant limitations in accuracy. When applied in combination to model ternary interactions, these methods often compound their inaccuracies, resulting in substantially reduced reliability. Moreover, tandem use fails to account for cooperative binding and the intricate conformational changes required, further diminishing their effectiveness.

State-of-the-art models, such as AlphaFold 3^3^-type models^4567^, represent a paradigm shift in ternary complex modeling. These models leverage end-to-end deep learning architectures capable of integrating multiple modalities, including protein sequences, DNA, RNA, and ligand information. This holistic approach allows for the prediction of complex biomolecular interactions with remarkable precision, capturing the intricate conformational change that traditional methods cannot. However, even with these advancements, these models still struggle to accurately model these molecular glue-induced interactions.

Here we present YDS-GlueFold (formerly known as YDS-Ternoplex), which was built upon commercially available AlphaFold 3-type models. Given AlphaFold 3’s demonstrated capabilities as a structure prediction foundation model, despite its limitations in handling molecular glue-induced ternary complexes accurately, we developed an enhanced approach. By incorporating guided diffusion into the AlphaFold 3-type model inference process, YDS-GlueFold effectively predicts molecular glue-induced ternary complex structures. We demonstrate YDS-GlueFold’s capabilities across eight diverse test cases of molecular glue-induced ternary complexes. Our approach accurately predicts structures for both E3 ligase-based systems (VHL:CDO1, CRBN with mTOR-FRB, NEK7, and VAV1-SH3c, and KBTBD4 with HDAC1) and non-E3 ligase complexes (FKBP12 with mTOR-FRB, BRD9 and QDPR), achieving RMSD values as low as 1.303 Å compared to experimental structures. Notably, YDS-GlueFold successfully predicts all seven novel protein-protein interfaces not present in training data. This capability is particularly evident in the FKBP12:MG:mTOR-FRB complex, where an alternative protein-protein interface exists in the training data that could potentially bias predictions toward this known interaction. YDS-GlueFold’s ability to correctly predict the novel interface instead of defaulting to the known interaction demonstrates its strong generalization capabilities rather than mere memorization of training data.

## Results

### Summary of Performance Metrics

To demonstrate the significant improvement YDS-GlueFold achieves over AlphaFold 3-type models, we compared our results with predictions from Chai-1r, an open-source implementation that closely replicates the AlphaFold 3 architecture.

**Table 1.** Summary of performance metrics across all test cases, comparing YDS-GlueFold and Chai-1r predictions against experimental structures.

| Case Complex |  | YDS-GlueFold |  |  | Chai-1r | Reference |
| --- | --- | --- | --- | --- | --- | --- |
|  |  | Best ipTM | RMSD (Å) | Correct Prediction | Correct Prediction |  |
| 1 | VHL:MG:CDO1 | 0.865 | 1.303 | ✓ | ✗ | PDB: 8VL9 |
| 2 | CRBN:MG:mTOR-FRB | 0.785 | N/A | ✓ | ✗ | Monte Rosa paper<sup>9</sup> |
| 3 | CRBN:MG:NEK7 | 0.797 | N/A | ✓ | ✗ | Monte Rosa paper<sup>9</sup> |
| 4 | CRBN:MG:VAV1-SH3c | 0.804 | N/A | ✓ | ✗ | Monte Rosa paper<sup>9</sup> |
| 5 | FKBP12:MG:mTOR-FRB | 0.848 | 1.462 | ✓ | X | PDB: 8PPZ |
| 6 | FKBP12:MG:BRD9 | 0.726 | N/A | ✓ | X | PDB: 9DU1* |
| 7 | FKBP12:MG:QDPR | 0.703 | N/A | ✓ | X | PDB: 9DTW* |
| 8 | KBTBD4:MG:HDAC1 | 0.816 | 2.263 | ✓ | X | PDB: 8VOJ |

\*Structure on hold for release
**Note:** The "Correct Prediction" columns indicate whether each model successfully predicted the experimental structure. For cases where quantitative RMSD assessment was not possible (N/A), we conducted detailed structural comparisons against the experimental analyses reported in the corresponding publications, validating that our predictions correctly captured the critical residue-specific interactions, binding motifs, and interface geometries described by the authors.

### Case 1: Ternary structure prediction of VHL:MG:CDO1

The first case is to predict the ternary complex of VHL:MG:CDO1, which has a solved structure in PDB (PDB ID: 8VL9). We specifically selected this case because no ternary complex structures of the VHL-CDO1 protein pair were included in the training set. This deliberate choice helps avoid suspicion of model memorization, making it an ideal case for evaluating the predictive capabilities of our method^8^.

Von Hippel-Lindau (VHL) functions as the substrate recognition component of the cullin-RING E3 ligase complex, playing a critical role in protein degradation via the ubiquitin-proteasome pathway. The protein of interest (POI) in this complex is cysteine dioxygenase type 1 (CDO1), an enzyme responsible for cysteine metabolism and regulation of oxidative stress. Dysregulation of CDO1 has been implicated in various cancers, particularly as a tumor suppressor. The biological significance of this case lies in demonstrating how molecular glues (MGs) can mediate interactions between metabolic enzymes like CDO1 and E3 ligases, potentially offering novel therapeutic strategies for targeting cancer-associated metabolic pathways^8^.

From our ensemble of sampled structures, we selected the model with the highest interface-predicted TM-score (ipTM), which reached 0.865. When compared to the experimentally determined experimental structure (PDB ID: 8VL9), this model showed excellent agreement with an RMSD of 1.303 Å. As shown in the structural alignment (Figure 1a), YDS-GlueFold effectively predicted the overall architecture of the ternary complex, and binding conformation of the MG.

**Figure 1.**
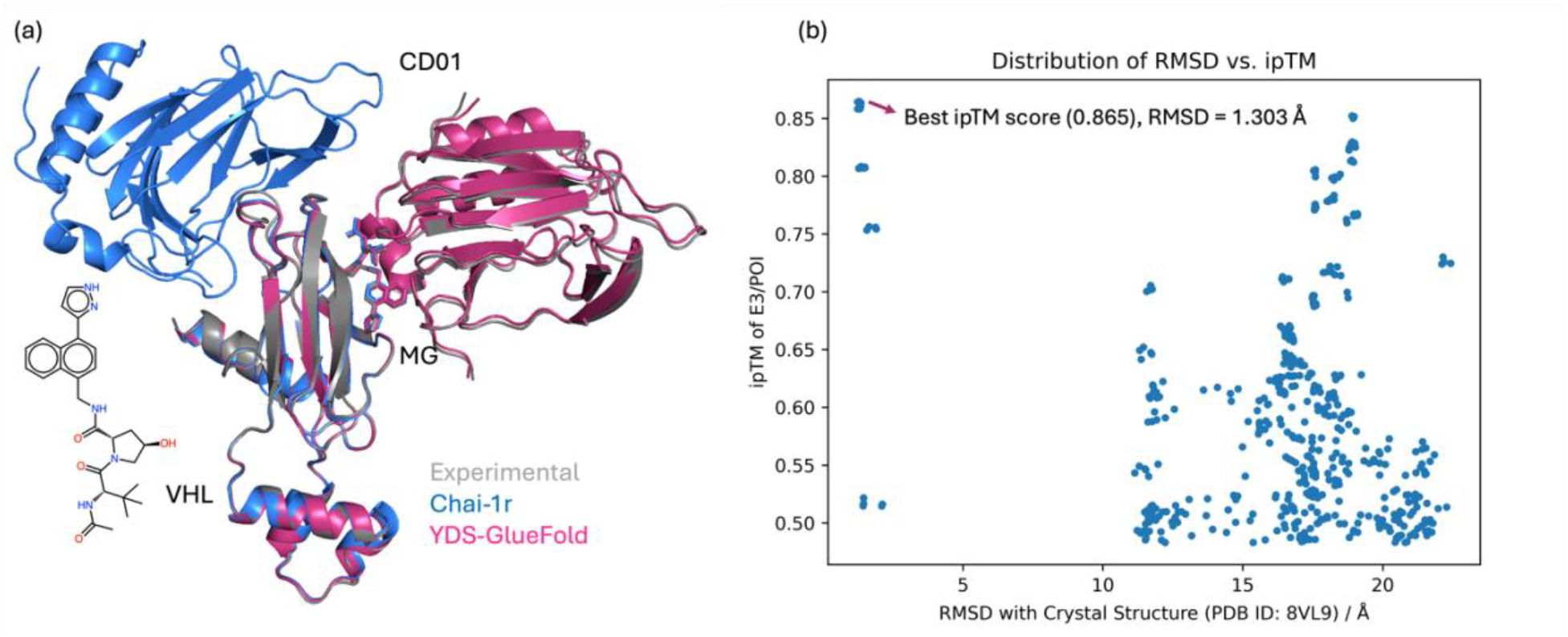
YDS-GlueFold prediction of the VHL:MG:CDO1 ternary complex. (a): Structural alignment of the experimental structure (gray), YDS-GlueFold prediction (pink), and Chai-1r prediction (blue). (b): Distribution of interface-predicted TM-score (ipTM) versus RMSD for YDS-GlueFold sampled structures (top 500) compared to the experimental structure (PDB ID: 8VL9). The best model, selected based on having the highest ipTM score (0.865) among all samples, shows an RMSD of 1.303 Å (indicated by arrow). RMSD was calculated using all heavy atoms.

**Figure 2.**
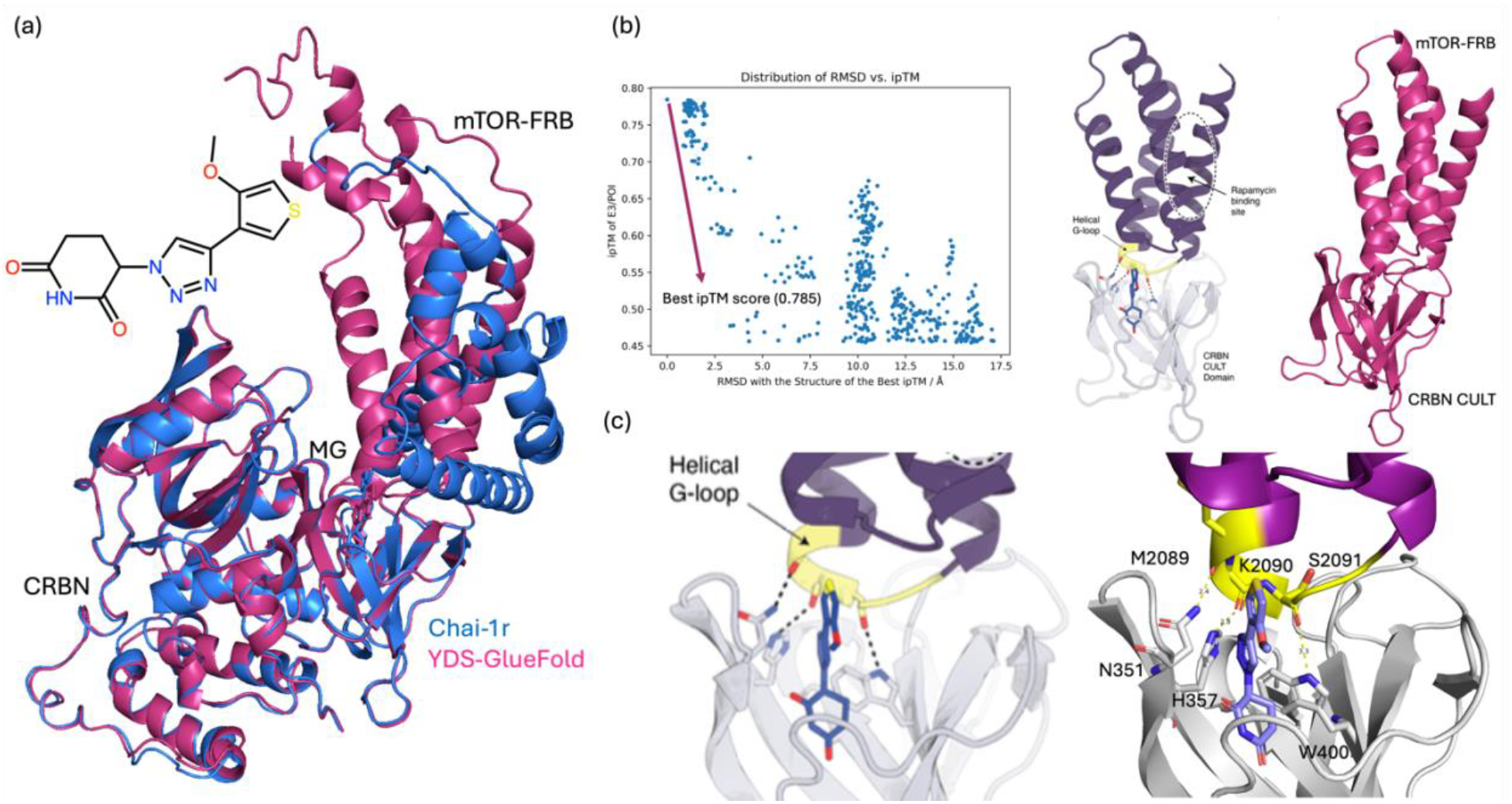
Predicted CRBN:MG:mTOR-FRB ternary complex and comparison with experimental analysis. (a) Overlay of predicted models YDS-GlueFold (pink) and Chai-1r (blue). (b) RMSD versus ipTM scatterplot of YDS-GlueFold sampled structures (top 500), with the highest scoring model (YDS-GlueFold) achieving an ipTM of 0.785. The reference structure for the RMSD calculation was the structure with the highest ipTM score. And separated views of CRBN CULT and mTOR-FRB domains, showing ligand-induced interaction regions. (c) Detailed view of the mTOR-FRB helical G-loop interacting with CRBN, showing ligand-induced stabilization through interactions involving key residues in mTOR-FRB (M2089, K2090, S2091) and CRBN (N351, H357, W400), consistent with experimental data.

**Figure 3.**
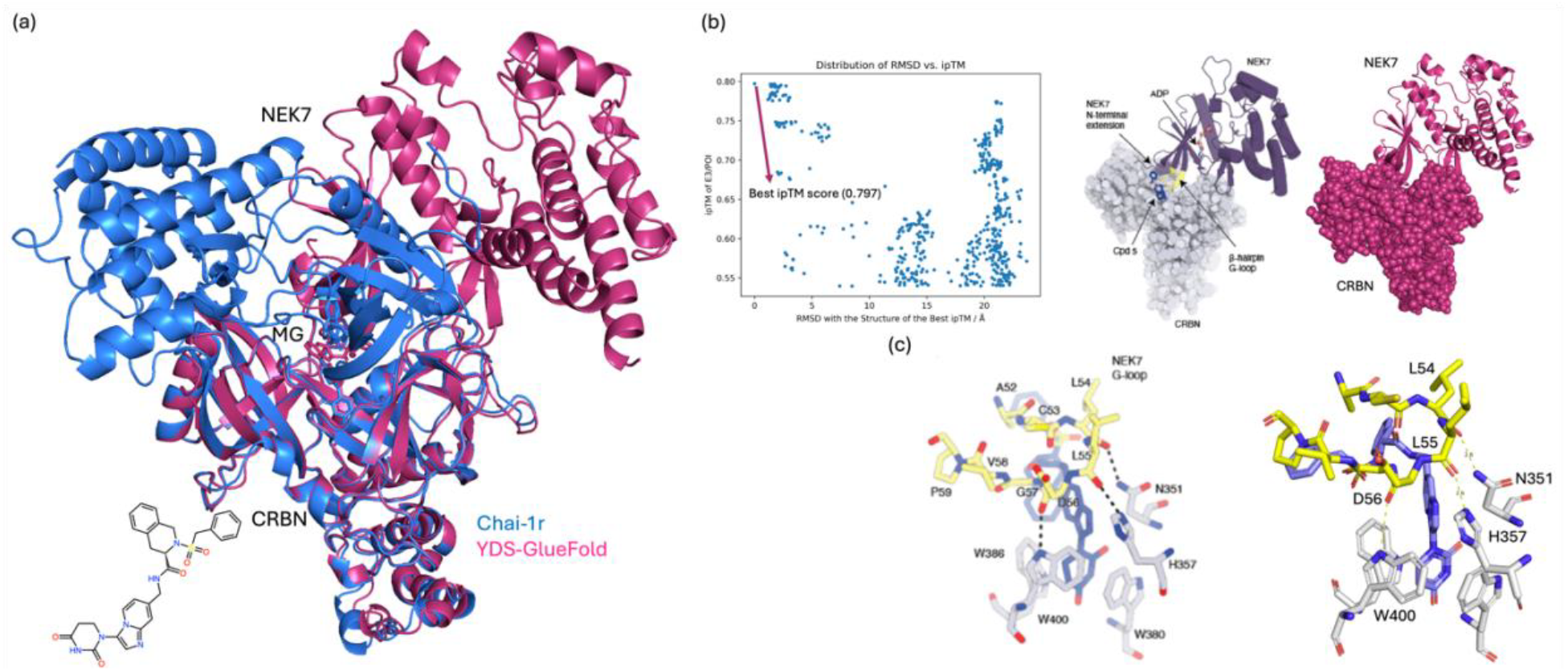
Predicted CRBN:MG:NEK7 ternary complex and comparison with experimental analysis. (a) Overlay of predicted models YDS-GlueFold (pink) and Chai-1r (blue). (b) RMSD versus ipTM scatterplot of YDS-GlueFold sampled structures (top 500), with the highest scoring model (YDS-GlueFold) achieving an ipTM of 0.797. The reference structure for the RMSD calculation was the structure with the highest ipTM score. Separated views of CRBN CULT and NEK7 domains highlight the ligand-induced interaction regions. (c) Detailed view of the NEK7 β-hairpin G-loop interacting with CRBN, showing ligand-induced stabilization through interactions involving key residues in NEK7 (L54, L55, D56) and CRBN (N351, H357, W400), consistent with experimental data.

### Case 2: Ternary structure prediction of CRBN:MG:mTOR-FRB

The second case explores the prediction of the ternary complex comprising CRBN, the mTOR-FRB domain, and a MG compound. mTOR (mammalian target of rapamycin) is a central regulator of cellular growth, metabolism, and survival, functioning as part of two distinct complexes, mTORC1 and mTORC2. Its FRB (FKBP12-rapamycin binding) domain plays a critical role in binding regulatory proteins and small molecules, making it a key target in therapeutic applications. CRBN (Cereblon) acts as the substrate receptor in the CRL4^CRBN^ E3 ubiquitin ligase complex, facilitating the degradation of neosubstrate proteins upon ternary complex formation^9^. The YDS-GlueFold prediction was selected from an ensemble of sampled structures with its highest inter-protein TM-score (ipTM) of 0.785.

Though there is no structure in PDB, recent research from Monte Rosa Therapeutics^9^ has disclosed a detailed analysis on their structure data. Analysis of the helical G-loop within the mTOR-FRB domain provided key insights into the ligand-induced stabilization of the ternary complex. The YDS-GlueFold model accurately captured critical interactions between mTOR-FRB residues (M2089, K2090, and S2091) and CRBN residues (N351, H357, and W400), forming a network of hydrogen bonds that align closely with experimentally validated structural data.

### Case 3: Ternary structure prediction of CRBN:MG:NEK7

This work presents the accurate prediction and experimental validation of the CRBN:NEK7 ternary complex facilitated by a molecular glue compound, emphasizing the reliability and consistency of the YDS-GlueFold model. NEK7 (NIMA-related kinase 7) is a serine/threonine kinase essential for microtubule organization and mitotic spindle assembly during cell division. Its dysregulation is associated with inflammatory responses and cancer, making it a target of interest^9^. The YDS-GlueFold prediction was selected from an ensemble of sampled structures with its highest inter-protein TM-score (ipTM) of 0.797. Though there is no structure in PDB, recent research from Monte Rosa Therapeutics^9^ has disclosed a detailed analysis on their structure data. The YDS-GlueFold model successfully predicted the key interactions between NEK7 residues (L54, L55, D56) and CRBN residues (N351, H357, and W400), forming a precise network of hydrogen bonds in strong agreement with experimental results.

### Case 4: Ternary structure prediction of CRBN:MG:VAV1-SH3c

In the fourth case, the POI is the SH3c domain of VAV1, a guanine nucleotide exchange factor (GEF) that plays a critical role in T-cell activation and cytoskeletal dynamics. The SH3c domain is crucial for mediating protein-protein interactions necessary for VAV1’s function in immune signaling. By forming a ternary complex with VAV1’s SH3c domain, molecular glues provide a unique mechanism to modulate immune pathways, which could have significant implications for immunotherapy^9^.

As illustrated by Figure 4, the structural prediction and experimental validation of the CRBN:VAV1 SH3c ternary complex induced by a molecular glue compound demonstrate the accuracy of the YDS-GlueFold model. The predicted structure with the highest inter-protein TM-score (ipTM) of 0.804, selected from an ensemble of sampled structures, shows a close match to experimentally validated binding poses. Though there is no structure in PDB, recent research from Monte Rosa Therapeutics^9^ has disclosed a detailed analysis on their structure data. Detailed views of YDS-GlueFold structure reveal that the VAV1 SH3c domain engages CRBN through the RT-loop, forming key hydrogen bonds between VAV1 residues (R796, D797, and S799) and CRBN residues (W400, H357, and N351). These interactions, as well as the molecular surface mimicry of the VAV1 RT-loop, are consistent with experimental observations, underscoring the predictive power of the YDS-GlueFold.

**Figure 4.**
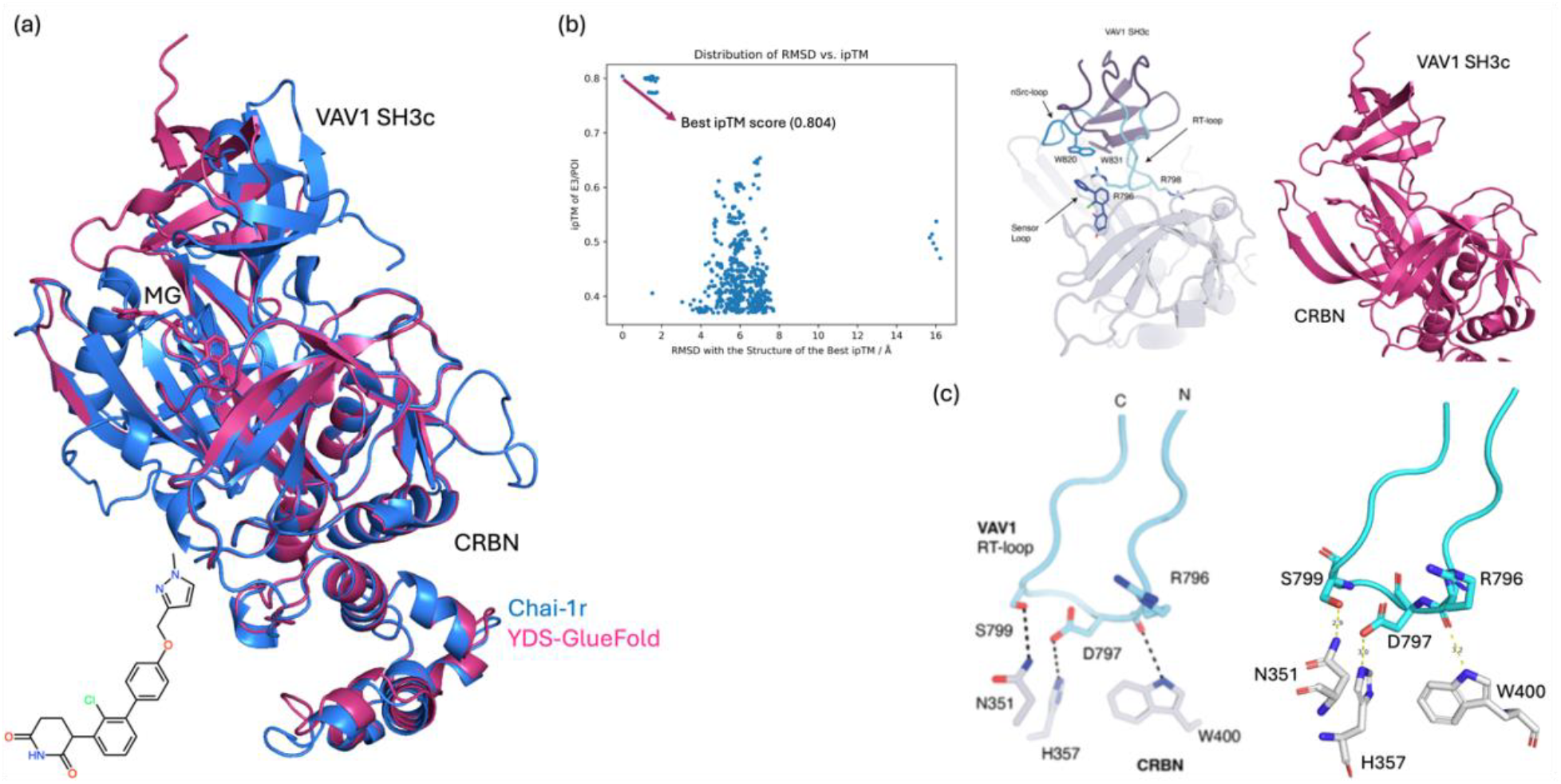
Predicted CRBN:MG:VAV1-SH3c ternary complex and comparison with experimental data. (a) Overlay of predicted models YDS-GlueFold (pink) and Chai-1r (blue). (b) RMSD versus ipTM scatterplot of YDS-GlueFold sampled structures (top 500), with the highest scoring model (YDS-GlueFold) achieving an ipTM of 0.804. The reference structure for the RMSD calculation was the structure with the highest ipTM score. Separated views of CRBN and VAV1-SH3c domains highlight the ligand-induced interaction regions. (c) Detailed view of the VAV1 RT-loop interacting with CRBN, showing ligand-induced stabilization through interactions involving key residues in VAV1 (R796, D797, and S799) and CRBN (W400, H357, and N351), consistent with experimental data.

### Case 5: Ternary structure prediction of FKBP12:MG:mTOR-FRB

The fifth case is the FKBP12:MG:mTOR-FRB ternary complex, which involves FKBP12 and the FRB domain of mTOR. Unlike the other cases, FKBP12 is not an E3 ligase but a prolyl isomerase that binds to immunosuppressive drugs such as rapamycin. There are a total of 11 PDB structures that include FKBP12, mTOR-FRB, and molecular glues. Remarkably, 10 of these structures share similar ternary configurations and protein-protein interfaces, whereas the most recent structure, PDB ID: 8PPZ, exhibits a completely different ternary structure and protein-protein interaction (PPI) pattern compared to the others^10^. Since 8PPZ was not included in the training set and the model never see this PPI during training, this case provides a unique opportunity to evaluate whether the model can correctly predict an unseen ternary configuration by learning potential atomic-level interactions.

As illustrated by Figure 5, the YDS-GlueFold model accurately predicts the FKBP12:MG:mTOR-FRB ternary complex. The structure with the highest inter-protein TM-score (ipTM) of 0.848 was selected from the ensemble of sampled models and aligns well with the experimental structure (PDB ID: 8PPZ), achieving a root-mean-square deviation (RMSD) of 1.462 Å. Without incorporating enhanced sampling inductive bias into the inference process, models like Chai-1r (blue) tend to predominantly sample protein-protein interactions (PPIs) that have been seen during training, as shown by the strong overlap of the Chai-1r prediction with the 1NSG structure. In our sampling, this type of PPI is associated with an ipTM around 0.789. The YDS-GlueFold model demonstrates its ability to generalize beyond seen PPIs by sampling and accurately predicting a novel ternary configuration, showcasing its capability to infer unseen PPIs and novel molecular glue-induced interactions.

**Figure 5.**
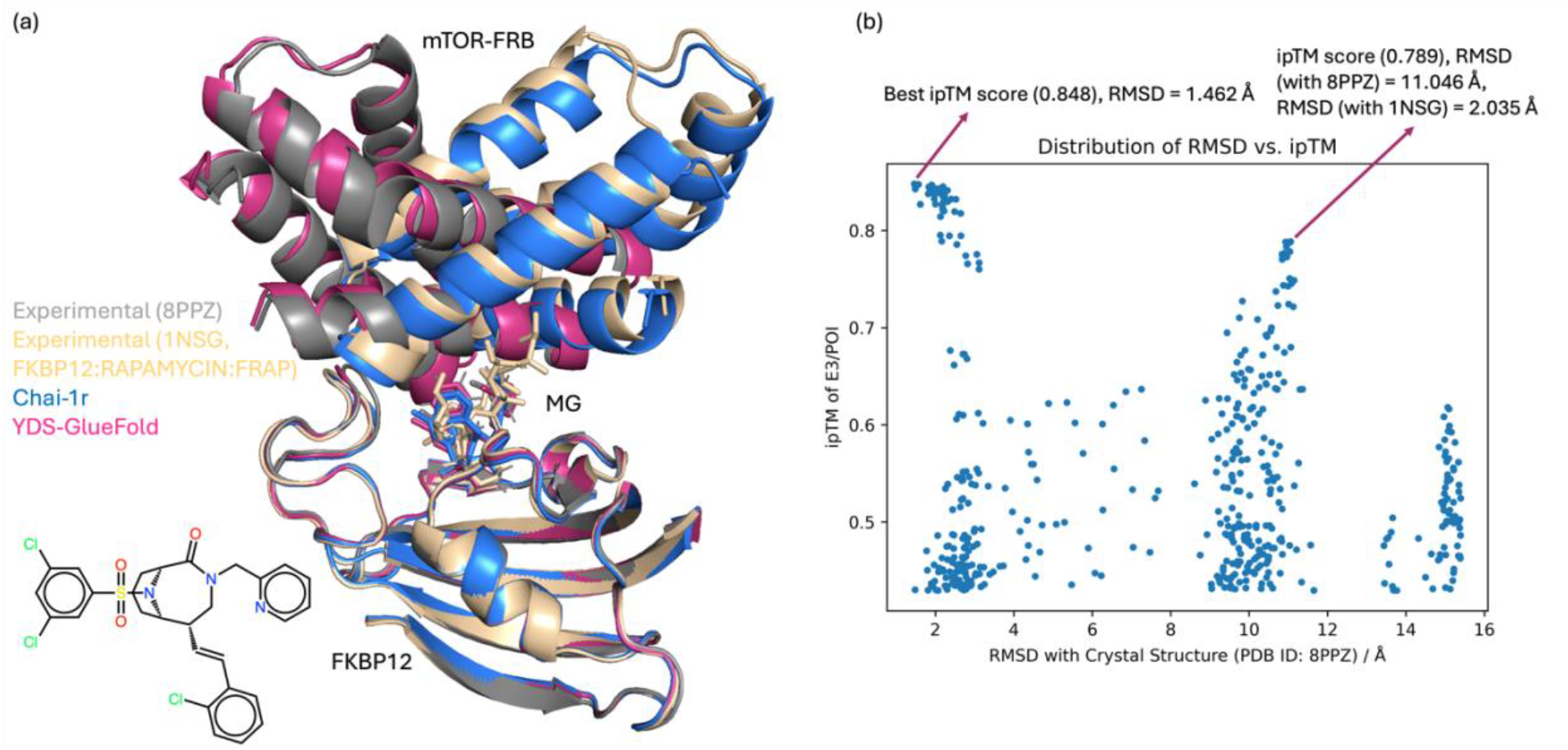
Predicted FKBP12:MG:mTOR-FRB ternary complex and comparison with experimental structures. (a) Overlay of the experimental structure (PDB ID: 8PPZ) with predicted models YDS-GlueFold (pink) and Chai-1r (blue), and another molecular glue rapamycin indued ternary structure 1NSG. (b) RMSD versus ipTM scatterplot of YDS-GlueFold sampled structures (top 500). The reference structure for RMSD calculation was 8PPZ.

**Figure 6.**
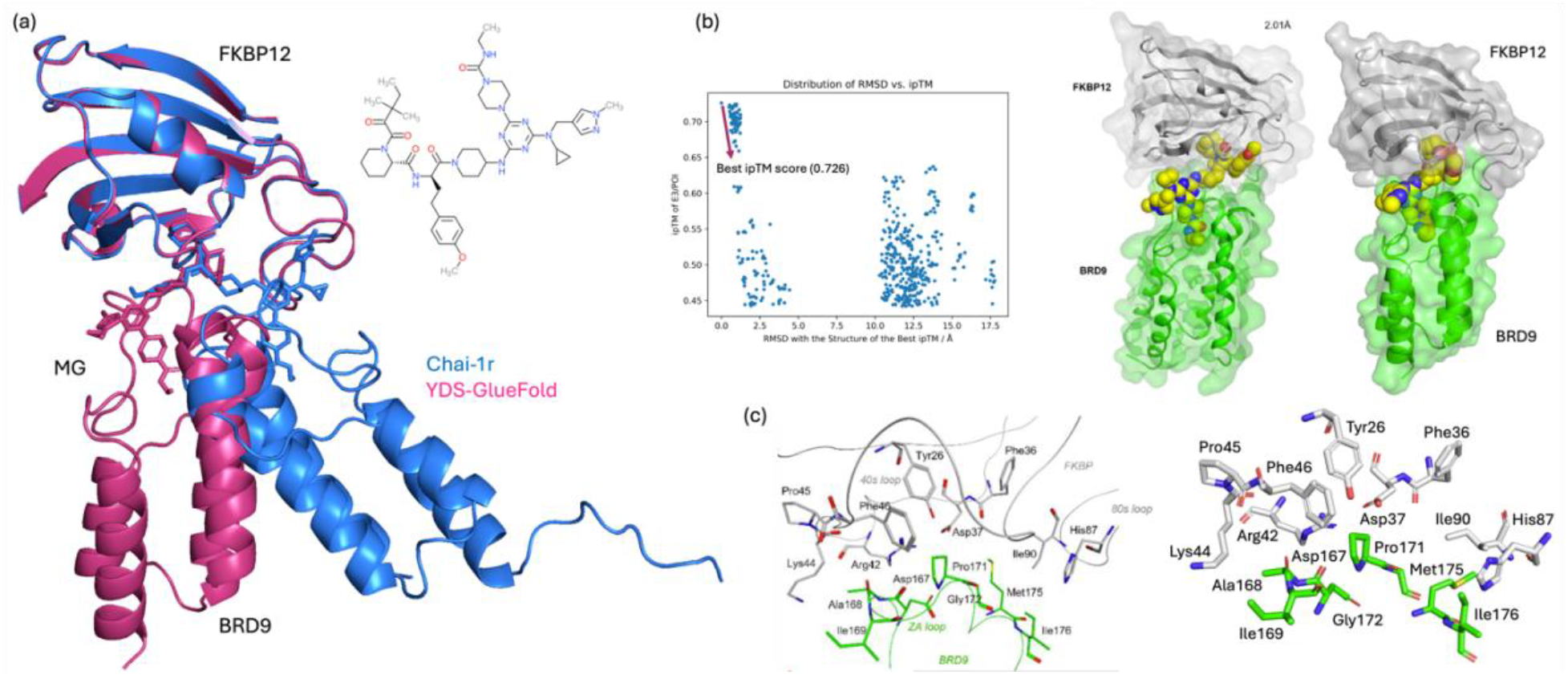
Predicted FKBP12:MG:BRD9 ternary complex and comparison with experimental analysis. (a) Overlay of predicted models YDS-GlueFold (pink) and Chai-1r (blue). (b) RMSD versus ipTM scatterplot of YDS-GlueFold sampled structures (top 500), with the highest scoring model (YDS-GlueFold) achieving an ipTM of 0.726. The reference structure for the RMSD calculation was the structure with the highest ipTM score. Separated views of FKBP12 and BRD9-Bromo domains, showing ligand-induced interaction regions. (c) Detailed view of the FKBP12 40s loop and 80s loop interacting with BRD9-Bromo ZA-loop, showing ligand-induced stabilization through interactions involving key residues in FKBP12 (I90) and BRD9-Bromo (M175, I176), consistent with experimental data.

**Figure 7.**
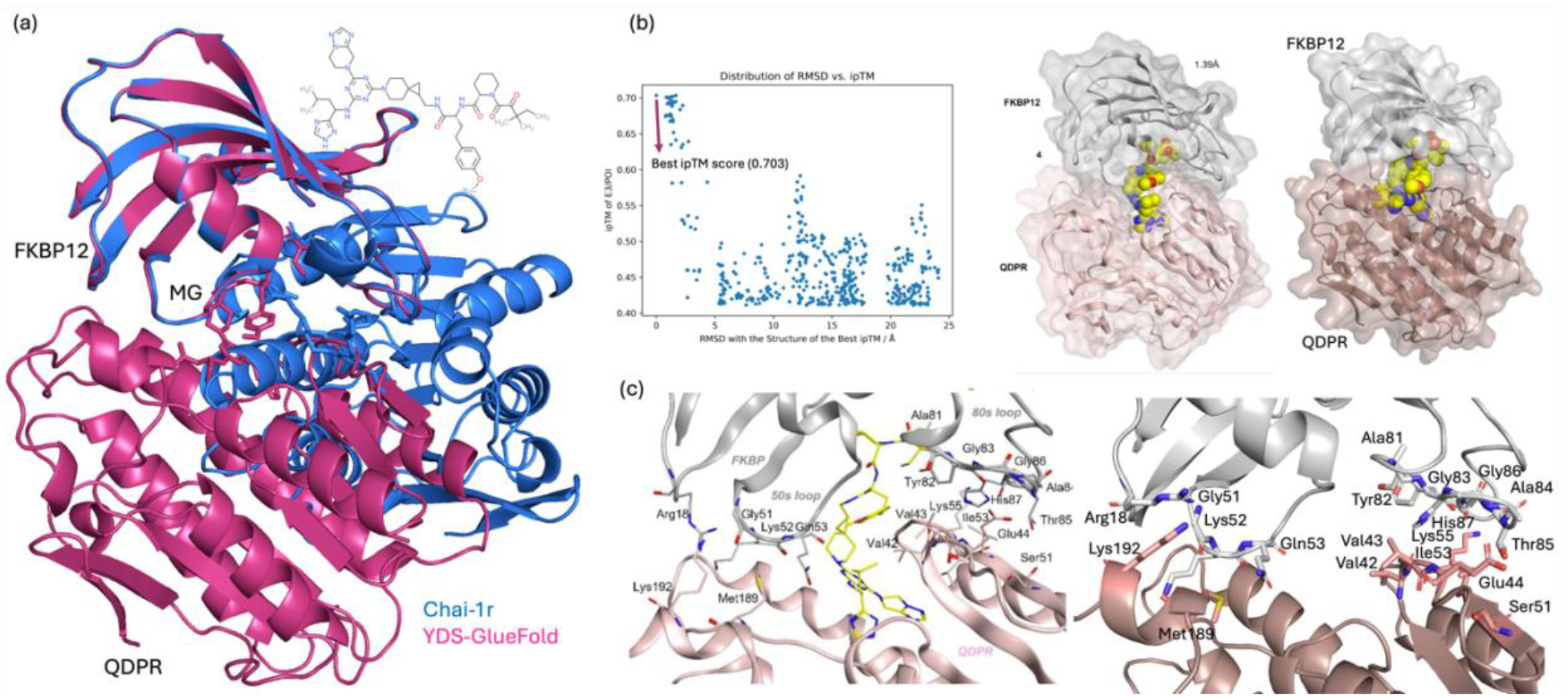
Predicted FKBP12:MG:QDPR ternary complex and comparison with experimental analysis. (a) Overlay of predicted models YDS-GlueFold (pink) and Chai-1r (blue). (b) RMSD versus ipTM scatterplot of YDS-GlueFold sampled structures (top 500), with the highest scoring model (YDS-GlueFold) achieving an ipTM of 0.703. The reference structure for the RMSD calculation was the structure with the highest ipTM score. Separated views of FKBP12 and QDPR, showing ligand-induced interaction regions. (c) Detailed view of the FKBP12 50s loop and 80s loop interacting with QDPR active site, showing ligand-induced stabilization through interactions involving key residues in FKBP12 (A84, T85) and QDPRo (E44), consistent with experimental data.

### Case 6: Ternary structure prediction of FKBP12:MG:BRD9

The sixth case investigates the FKBP12:MG:BRD9 ternary complex. A detailed structural analysis disclosed in a bioRxiv paper by researchers at Novartis Biomedical Research. The PDB structure (PDB ID: 9DU1) remains on hold for release^12^. Similar to previous cases, this system was deliberately selected because no structures of the FKBP12-BRD9 protein pair were included in the training set, ensuring that the YDS-GlueFold model generalizes beyond training data used for train Alphafol3-type models.

In this complex, FKBP12 remains the adapter protein, and serves as a key adapter to recruit BRD9, a bromodomain-containing protein that plays a pivotal role in chromatin remodeling and gene transcription regulation. BRD9 is a component of the non-canonical BAF (ncBAF) chromatin remodeling complex, and its dysregulation has been implicated in cancer, particularly synovial sarcoma and other malignancies, making it a promising therapeutic target^13^. The ability of molecular glues to mediate interactions between FKBP12 and BRD9 showcases their versatility in targeting chromatin-associated proteins. This case demonstrates how molecular glues can facilitate novel protein-protein interactions to modulate epigenetic regulators, offering new therapeutic avenues for cancer and other diseases linked to chromatin dysregulation.

The YDS-GlueFold model predicted the ternary structure with a highest interface-predicted TM-score (ipTM) of 0.726. Analysis of the predicted structure showed excellent agreement with the bioRxiv analysis, particularly regarding the ZA-loop of the bromodomain. Key hydrophobic interactions were accurately captured, including those between Met175 and Ile176 of BRD9 and Ile90 of the FKBP12 80s loop. Additional protein-protein interactions between the N-terminal portion of the ZA loop and the 40s loop of FKBP12 further highlighted the model’s ability to predict critical interactions^12^.

### Case 7: Ternary structure prediction of FKBP12:MG:QDPR

The seventh case is the FKBP12:MG:QDPR ternary complex. Structural details also disclosed in the bioRxiv publication by Novartis Biomedical Research. The associated PDB structure (PDB ID: 9DTW) remains on hold for release^12^. Like the FKBP12:BRD9 complex, this case was selected because no FKBP12-QDPR protein pair structures were included in the training data, making it a stringent test of the model’s predictive capabilities.

In this system, FKBP12 remains the adapter protein, while QDPR (quinoid dihydropteridine reductase) is the protein of interest. QDPR is an enzyme involved in the regeneration of tetrahydrobiopterin (BH4), an essential cofactor for several critical enzymatic reactions, including neurotransmitter synthesis^14^. Dysregulation of QDPR has been associated with hyperphenylalaninemia and neurological disorders, making it an important therapeutic target. The molecular glue mediates the recruitment of QDPR by FKBP12, forming a ternary complex that highlights the potential of this approach in targeting enzymes related to metabolic and neurological disorders. This case underscores the capability of molecular glues to exploit non-canonical interactions to address diseases linked to metabolic dysfunctions and enzymatic dysregulation.

The YDS-GlueFold model sampled a highest ipTM of 0.703 in predicting the FKBP12:MG:QDPR complex. Structural analysis revealed precise modeling of interactions between the 50s and 80s loops of FKBP12 and the regions flanking QDPR’s active site. Notably, hydrogen bonding interactions between FKBP12 residues Ala84 and Thr85 and QDPR residue Glu44 were faithfully reproduced, demonstrating the model’s ability to capture atomic-level details.

### Case 8: Ternary structure prediction of KBTBD4:MG:HDAC1

The eighth case examines the KBTBD4:MG:HDAC1 ternary complex, for which a high-resolution structure has been determined (PDB ID: 8VOJ)^15^. KBTBD4 is a BTB (Broad-Complex, Tramtrack, and Bric-a-brac) domain-containing substrate adaptor protein for the CUL3-RING E3 ligase complex. It functions by recruiting target proteins to the ligase complex for ubiquitination and subsequent degradation. HDAC1 (histone deacetylase 1), the protein of interest in this ternary complex, is a critical epigenetic regulator that modulates gene expression by removing acetyl groups from histone tails. HDAC1 is implicated in numerous cellular processes, including cell cycle progression, differentiation, and oncogenesis, making it a compelling therapeutic target in cancer and neurological disorders. Importantly, no structural data of the KBTBD4-HDAC1 protein pair was included in the training set, ensuring that this system serves as an independent and unbiased test of the model’s generalization ability^16,17^.

From the ensemble of sampled structures, YDS-GlueFold predicted a ternary configuration with a highest ipTM of 0.816, indicating high confidence in the modeled protein-protein interface. As illustrated in Figure 8, the YDS-GlueFold model accurately recapitulates the KBTBD4:MG:HDAC1 ternary complex, showing strong agreement with the experimental structure (PDB ID: 8VOJ) and achieving an RMSD of 2.263 Å. Structural analysis revealed that the model precisely captures key hydrophobic interactions—specifically, Ile310 and Leu356 of KBTBD4-B form direct contacts with Phe150 and Phe205 of HDAC1, respectively. These residues surround the molecular glue and nucleate a hydrophobic core at the protein-protein interface, stabilizing the ternary assembly^15-17^.

**Figure 8.**
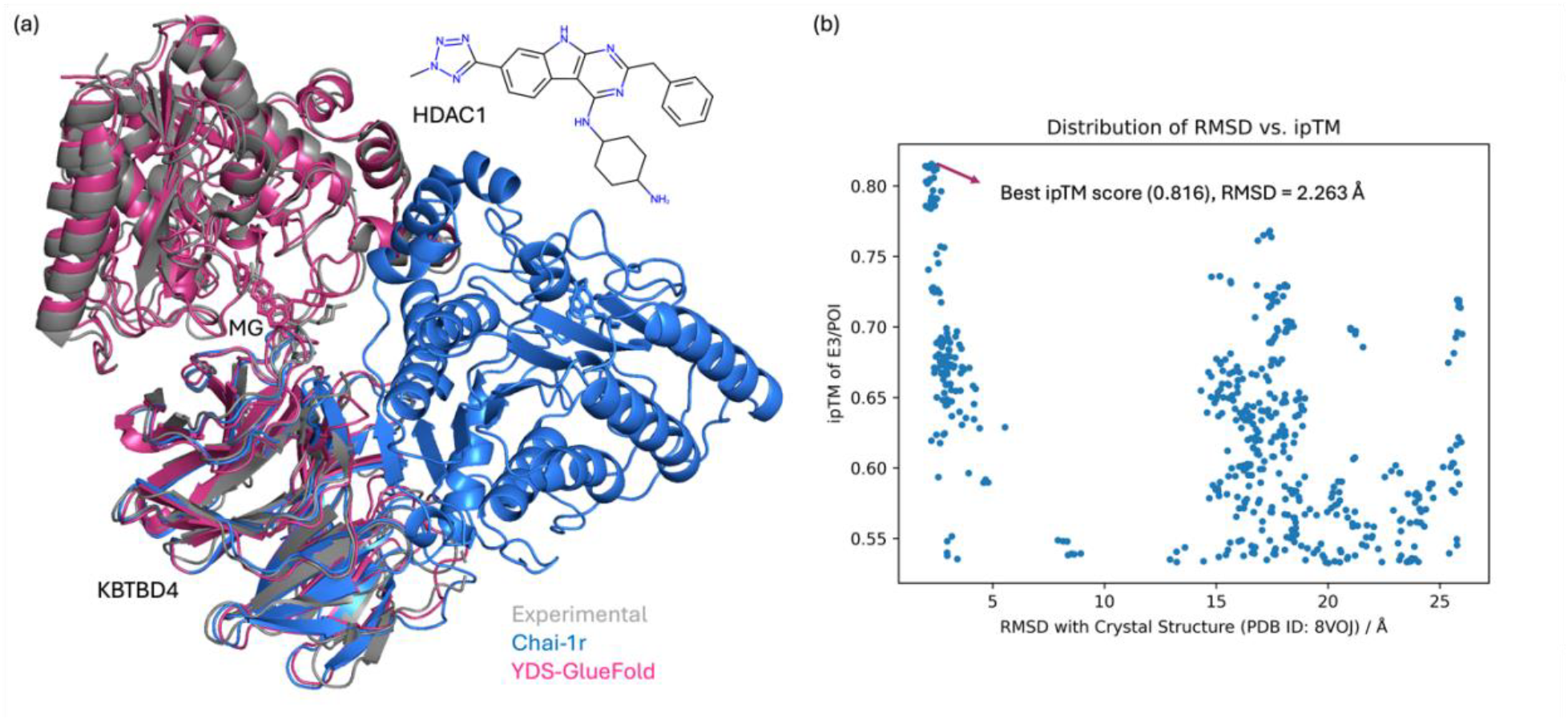

The KBTBD4:MG:HDAC1 complex illustrates how molecular glues can reprogram substrate specificity of E3 ligases to redirect the degradation machinery toward disease-relevant epigenetic regulators. This case further demonstrates the capacity of YDS-GlueFold to correctly predict novel ternary architectures, including previously unseen protein-protein interfaces, by leveraging atomic-level interaction modeling rather than memorization of training data.

## Discussion

The YDS-GlueFold model demonstrates substantial advancements in predicting molecular glue-induced ternary complexes by incorporating guided diffusion modules. By building on AlphaFold 3-type models, YDS-GlueFold not only achieves higher predictive accuracy but also overcomes the limitations of existing approaches that tend to overfit to seen protein-protein interaction patterns or fail to capture cooperative binding central to molecular glue mechanisms.

Our results highlight the model’s capability to generalize across diverse ternary complexes, including both E3 ligase-based (e.g., VHL:CDO1, CRBN:NEK7) and non-E3 ligase systems (e.g., FKBP12:mTOR-FRB). In each case, the model accurately captured critical interaction interfaces and ligand-induced stabilization, evidenced by low RMSD values and strong alignment with experimental data. Notably, the FKBP12:mTOR-FRB case exemplifies YDS-GlueFold’s ability to infer entirely novel configurations, distinguishing it from baseline models that default to memorized interaction patterns.

Among our test cases, Case 8 (KBTBD4:MG:HDAC1) presented the most significant predictive challenge, followed by Case 4 (CRBN:MG:VAV1-SH3c). This difficulty can be attributed to the presence of flexible loop regions in the interaction interfaces. Loops inherently exhibit greater conformational variability than structured elements, necessitating accurate sampling of their possible orientations to correctly identify the binding interface. The ability of YDS-GlueFold to accurately predict these challenging interface configurations without prior knowledge of interaction sites demonstrates the effectiveness of our guided diffusion approach in efficiently exploring complex conformational landscapes.

The success of YDS-GlueFold in achieving state-of-the-art performance by adding guided diffusion modules exclusively during the inference process underscores a significant implication: the Pairformer module has effectively learned the interaction landscape. This suggests that the underlying latent space is adept at capturing the foundational features of high dimensional interactions enabling robust predictions with only inference-stage adjustments. This efficiency not only validates the design of the Pairformer architecture but also highlights its potential for broader applications in mapping interaction to function.

The integration of guided diffusion is central to this success. By drawing on principles from statistical mechanics and diffusion processes, the model efficiently navigates high-dimensional conformational landscapes, overcoming energy barriers that limit traditional sampling approaches. This targeted exploration of rare but biologically significant configurations ensures robust predictions, even for complexes with limited structural precedence.

Moreover, the theoretical framework of inference scaling laws^11^ underpins our approach, emphasizing that indiscriminate increases in computational resources do not guarantee performance gains. Instead, strategic enhancements to the inference process, such as our inductive bias mechanism, drive meaningful improvements. This insight underscores the importance of pairing computational resources with methodological innovations to achieve scalable and efficient predictions.

In conclusion, YDS-GlueFold represents a significant leap forward in computational modeling of molecular glue-induced ternary complexes. By addressing the intricate interplay of cooperative binding, it opens new avenues for rational drug design targeting previously undruggable proteins. These advancements pave the way for more precise and effective therapeutic strategies, expanding the scope of molecular glue applications in drug discovery.

## Supporting information

Case 8 KBTBD4:MG:HDAC1

## Reference

1. Garber, K. The glue degraders. Nat Biotechnol 42, 546–550 (2024).

2. Konstantinidou, M. & Arkin, M. R. Molecular glues for protein-protein interactions: Progressing toward a new dream. Cell Chemical Biology 31, 1064–1088 (2024).

3. Abramson, J. et al. Accurate structure prediction of biomolecular interactions with AlphaFold 3. Nature 630, 493–500 (2024).

4. Wohlwend, J. et al. Boltz-1 Democratizing Biomolecular Interaction Modeling. Preprint at 10.1101/2024.11.19.624167 (2024).

5. Chai Discovery et al. Chai-1: Decoding the molecular interactions of life. Preprint at 10.1101/2024.10.10.615955 (2024).

6. Qiao, Z. et al. NeuralPLexer3: Accurate Biomolecular Complex Structure Prediction with Flow Models. Preprint at 10.48550/arXiv.2412.10743 (2024).

7. Protenix_Technical_Report.

8. Tutter, A. et al. A small molecule VHL molecular glue degrader for cysteine dioxygenase 1. Preprint at 10.1101/2024.01.25.576086 (2024).

9. Petzold, G. et al. Mining the CRBN Target Space Redefines Rules for Molecular Glue-induced Neosubstrate Recognition. Preprint at 10.1101/2024.10.07.616933 (2024).

10. Deutscher, R. C. E. et al. Discovery of fully synthetic FKBP12-mTOR molecular glues. Preprint at 10.26434/chemrxiv-2023-4vb0m-v2 (2024).

11. Wu, Y., Sun, Z., Li, S., Welleck, S. & Yang, Y. Inference Scaling Laws: An Empirical Analysis of Compute-Optimal Inference for Problem-Solving with Language Models. Preprint at 10.48550/arXiv.2408.00724 (2024).

12. Zandi, T. A. et al. Discovery of molecular glues that bind FKBP12 and novel targets using DNA-barcoded libraries. Preprint at 10.1101/2024.12.09.627499 (2024).

13. Helming, K. C., Wang, X., Roberts, C. W. M. Vulnerabilities of mutant SWI/SNF complexes in Cancer. Cancer Cell 26, 309–317 (2014).

14. Thony, B., Auerbach, G., Blau, N. Tetrahydrobiopterin biosynthesis, regeneration and functions. Biochemical Journal 347, 1–16 (2000).

15. Megan J.R. Yeo. et al. Asymmetric Engagement of Dimeric CRL3KBTBD4 by the Molecular Glue UM171 Licenses Degradation of HDAC1/2 Complexes. Preprint at 10.1101/2024.05.14.593897 (2024).

16. Megan J.R. Yeo. et al. UM171 glues asymmetric CRL3–HDAC1/2 assembly to degrade CoREST corepressors. Nature 639, 232–240 (2025).

17. Xie, X. et al. Converging mechanism of UM171 and KBTBD4 neomorphic cancer mutations. Nature 639, 241–249 (2025).

